# Enhanced Astrocyte Responses are Driven by a Genetic Risk Allele Associated with Multiple Sclerosis

**DOI:** 10.1101/206110

**Authors:** Gerald Ponath, Matthew R. Lincoln, Somiah Dahlawi, Mayyan Mubarak, Tomokazu Sumida, Laura Airas, Shun Zhang, Cigdem Isitan, Thanh D. Nguyen, Cedric S. Raine, David A. Hafler, David Pitt

## Abstract

Epigenetic annotation studies of genetic risk variants for multiple sclerosis (MS) implicate dysfunctional lymphocytes in MS susceptibility; however, the role of central nervous system (CNS) cells remains unclear. We investigated the effect of the risk variant, rs7665090^G^, located near *NFKB1*, on astrocytes. We demonstrated that chromatin is accessible at the risk locus, a prerequisite for its impact on astroglial function. The risk variant was associated with increased NF-κB signaling and target gene expression driving lymphocyte recruitment in cultured human astrocytes and astrocytes within MS lesions, and with increased lesional lymphocytic infiltrates. In MS patients, the risk genotype was associated with increased lesion volumes on MRI. Thus, we established that the rs7665090^G^ variant perturbs astrocyte function resulting in increased CNS access for peripheral immune cells. MS may thus result from variant-driven dysregulation of the peripheral immune system and the CNS, where perturbed CNS cell function aids in establishing local autoimmune inflammation.

**One Sentence Summary:** The NF-κB relevant multiple sclerosis risk variant, rs7665090^G^, drives astrocyte responses that promote lesion formation.

## Introduction

Multiple sclerosis (MS) is a genetically mediated inflammatory disease of the central nervous system (CNS), where infiltrating immune cells lead to focal, demyelinating lesions^1^. Genome-wide association studies (GWAS) have now identified over 200 genetic variants that confer increased risk of developing MS^2^. A recent epigenetic annotation study has demonstrated that MS risk variants are highly enriched in immune enhancers that are active in T and B cells^3^, suggesting that risk variant-mediated MS susceptibility is driven by changes in gene regulation in lymphocytes. This was confirmed in our recent work where we prioritized up to 551 potentially associated MS susceptibility genes and found that they implicated multiple innate and adaptive pathways distributed across different cells of the immune system^4^. It remains unclear whether genetic variants only affect immune cells or also change CNS cell function, thereby driving MS risk by altering CNS-intrinsic pathways.

We addressed this question by investigating how a common MS risk variant, rs7665090^G^, drives NF-κB signaling in astrocytes. This risk variant has recently been shown to increase NF-κB p50 expression and NF-κB activation in lymphocytes^5^. As a master regulator of the innate and adaptive immunity, NF-κB plays a critical role in autoimmunity, including MS, where 18% of all allelic MS risk variants are estimated to affect the NF-κB signaling pathway^5,6^.

NF-κB signaling also plays a role in activation of astrocytes, a cell type that is critically involved in the formation of MS white matter lesions. Astrocytes help limit the entry of peripheral immune cells into the CNS by forming a physical barrier, glia limitans, but can also mount powerful pro-inflammatory responses, including expression of adhesion molecules and secretion of chemokines that lead to leukocytes recruitment^7^. Astrocyte-specific inhibition of NF-κB activation has been shown to dramatically ameliorate immune infiltration and tissue damage in experimental autoimmune encephalomyelitis (EAE), an animal model of MS^8,9^, suggesting that genetic variants relevant to NF-κB may alter astrocyte responses and astrocyte-mediated MS lesion pathology.

The rs7665090 risk variant tags a haplotype block of more than 90 variants that are in tight linkage disequilibrium. The haplotype block spans a large number of gene regulatory elements and lies within *MANBA* and the intergenic space between *NFKB1* and *MANBA* as mapped by the ENCODE project (Fig. 1)^5,10,11^. The rs7665090^G^ variant has an approximate frequency of 55% in the general population and increases the odds ratio for MS susceptibility by 1.09 per G allele carried^6^.

**Figure 1.**
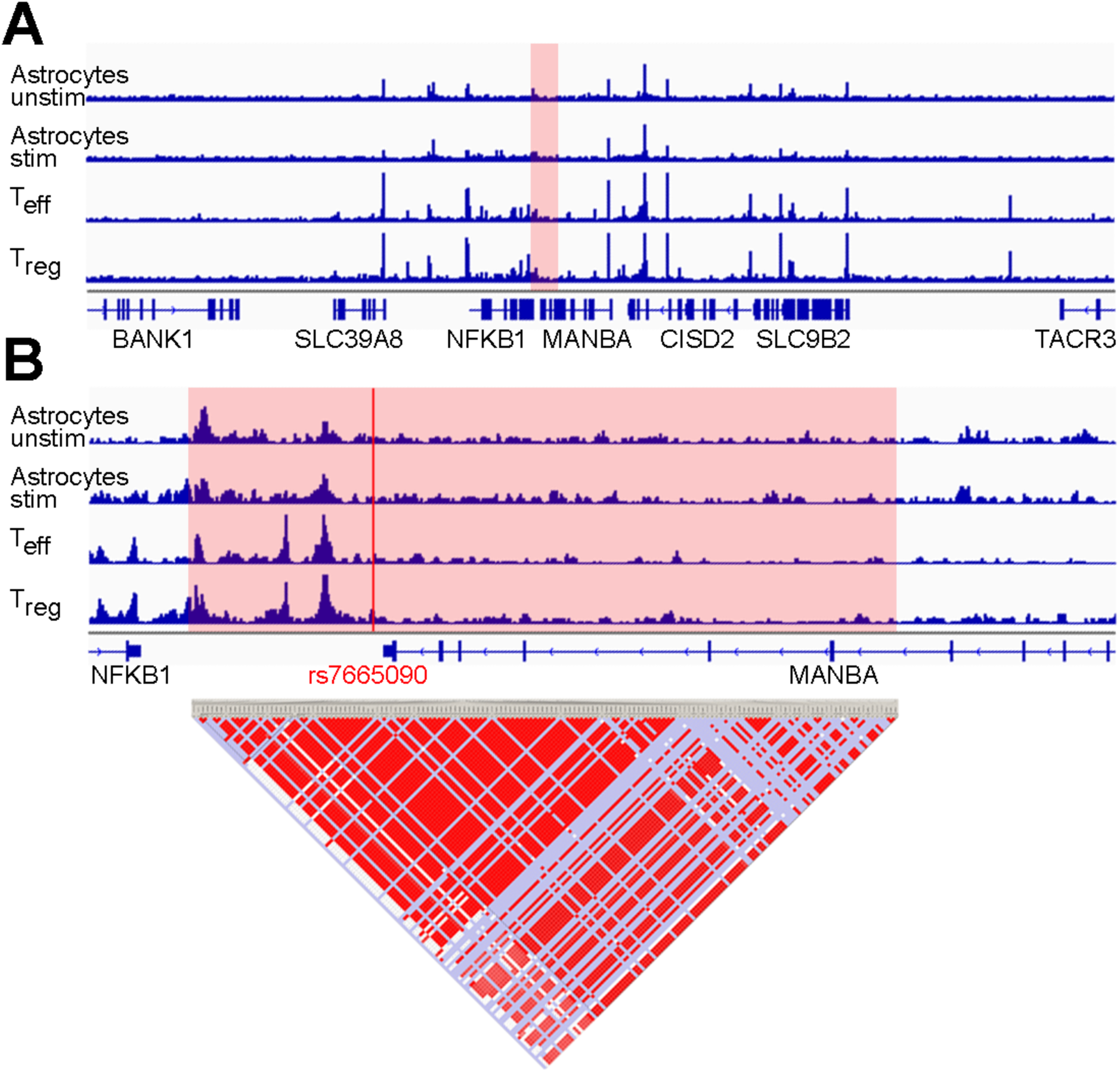
Chromatin accessibility in the *NFKB1/MANBA* risk haplotype block in astrocytes and T cells. (**A**) Normalized ATAC-seq profiles in the *NFKB1/MANBA* risk locus are similar in unstimulated and stimulated human fetal astrocytes (AC_unstim_ and AC_stim_; first two tracks) and in *ex vivo* unstimulated effector and regulatory T cells (T_eff_ and T_reg_; third and fourth track). The height of the bar graph at each point represents the number of unique, singly-mapping reads at each genomic position; each track is normalized for library size. The *NFKB1/MANBA* haploblock is shaded in red. The genomic coordinates are chr4:101,796383-103694,281. (**B**) Higher magnification (chr4:102,614,123-102,674,492) of the risk haploblock, shaded in red, shows accessible chromatin mostly in the intergenic region between *NFKB1* and *MANBA* with comparable ATAC-seq profiles in astrocytes and T cells. The location of the tagging SNP rs7665090 is indicated by the red line. Pairwise linkage disequilibrium (D’) values between SNPs in the CEU population of the 1000 Genomes Project are indicated below (red indicates D’>0.95).

## Results

### Chromatin accessibility in the *NFKB1/MANBA* risk haplotype block in astrocytes and T cells

We assessed chromatin accessibility within the *NFκB1*-*MANBA* risk haplotype block tagged by rs7665090 with an assay for transposase-accessible chromatin with high throughput sequencing (ATAC-seq) on FACS-sorted unstimulated and stimulated human fetal and iPSC-derived astrocytes and for comparison, on *ex vivo*, unstimulated human effector and regulatory T cells^12^. This haplotype block spans the intergenic region downstream of the *NFKB1* gene and a part of the MANBA gene. Open chromatin sites captured by ATAC-seq were restricted to the intergenic region and highly congruent between both astrocytes and T cells (fig.1), suggesting that the risk haplotype block-associated increase in *NFKB1* gene expression shown previously in T cells^5^, may also apply to astrocytes. In contrast, chromatin at a risk locus within the intragenic region of *NFKB1* that is associated with non-CNS autoimmune diseases (inflammatory bowel disease and systemic scleroderma), was accessible only in T cells but not in astrocytes (Fig. S1). The specificity of our data set was further confirmed by the mutually exclusive chromatin accessibility in lineage-specific genes of astrocytes and T cells (GFAP, CD2). Thus, our results indicate that while *NFKB1* is expressed by a multitude of cell types, accessibility of different regulatory enhancer regions for the *NFKB1* gene vary among different cell types. As expected, chromatin accessibility was not affected by the rs7665090 genotype, i.e. did not differ between astrocytes or lymphocytes with the risk and protective genotype (data not shown).

### Effect of the rs7665090^G^ variant on activated human iPSC-derived astrocytes

Next, we determined the impact of the rs7665090 risk variant on NF-κB signaling in iPSC-derived astrocytes generated from fibroblasts of MS patients that are homozygous either for the risk (rs7665090^GG^) or protective variant (rs7665090^AA^) (Fig. S7). In unstimulated astrocytes, NF-κB signaling was low in both groups, as measured by degradation of inhibitor of NF-κBα (IκBα) and phosphorylation of p65. Treatment of astrocytes with a combination of TNFα and IL-1β, cytokines that induce or enhance NF-κB signaling and play a major role during MS lesion development^13–15^, increased NF-κB activation in both groups; however, NF-κB activation was significantly higher in astrocytes carrying the risk variant (Fig. 2, A and B). In addition, at a resting state, expression of NF-κB p50 but not of p65 was higher in astrocytes with the risk variant compared to astrocytes with the protective variant. After stimulation, both p50 and p65 expression was substantially upregulated, again significantly higher in astrocytes with the risk variant (Fig. 2C).

**Figure 2.**
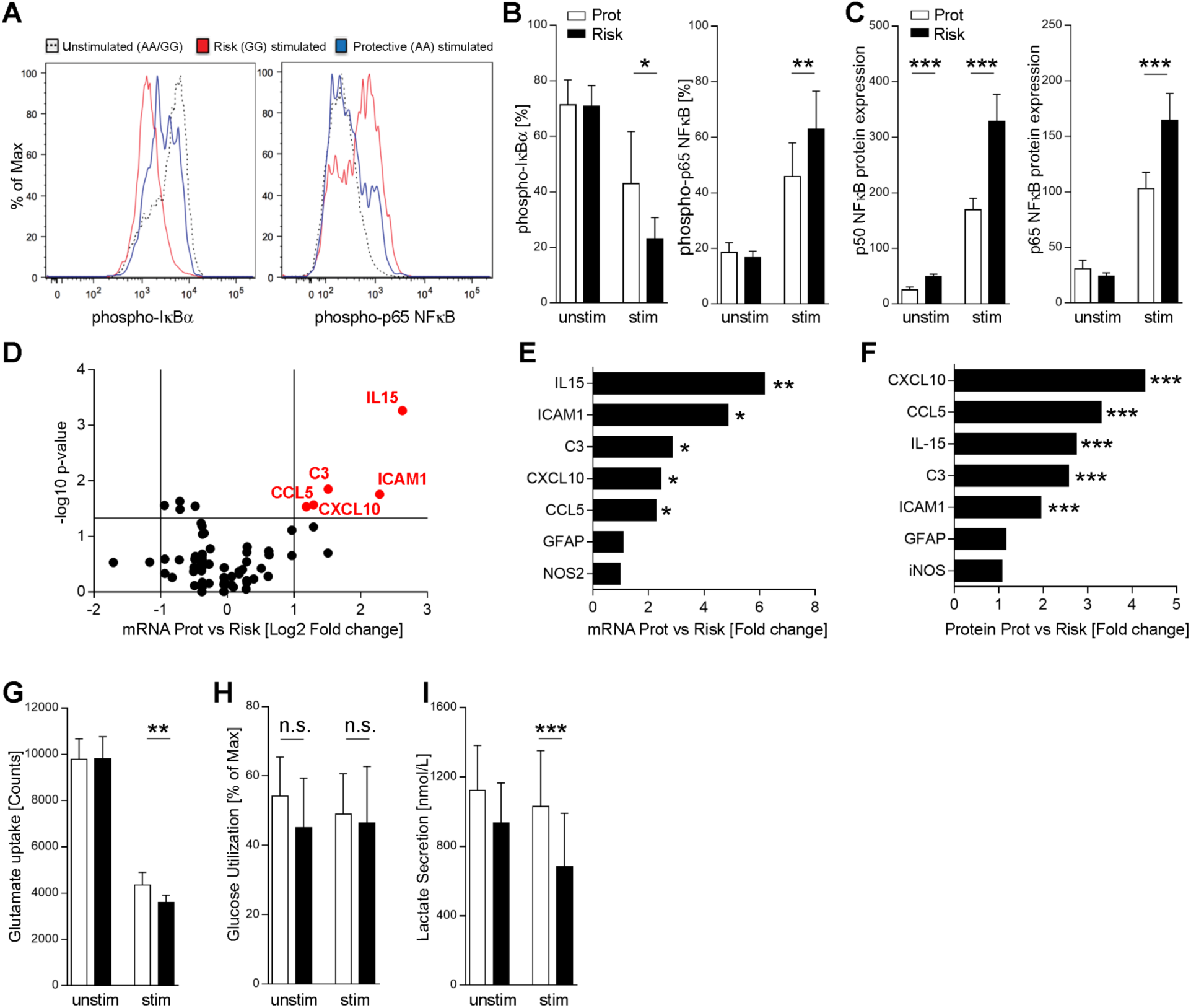
Effect of the rs7665090^G^ variant on activated human iPSC-derived astrocytes. (**A**, **B**) Degradation of IκBα and phosphorylation of p65 in iPSC-derived astrocytes with risk and protective variant at a resting state and after 10 min of stimulation with TNFα (50 ng/ml) and IL-1β (10 ng/ml) by flow cytometry. (**C**) Expression of p50 and p65 in unstimulated and stimulated (10 ng/ml IL-1β; 100 U/ml IFNγ for 12 hrs, followed by 50 ng/ml TNFα for 4 hrs) iPSC-derived astrocytes from both groups by Western blot. (**D**-**E**) Volcano plot profiling astroglial expression of 84 *NFKB* target genes after stimulation with IL-1β, IFNγ and TNFα. Red dots indicate genes with ≥2-fold expression and FDR-adjusted *p*-values ≤0.05. Genes with significantly increased expression are listed in (**E**) and corresponding protein expression in (**F**). (**G**) Uptake of L-[3,4-^3^H] glutamic acid by iPSC-derived unstimulated and stimulated (with IL-1β, IFNγ and TNFα) astrocytes with risk and protective variant. (**H**, **I**) Glucose uptake and lactate secretion by astrocytes with the risk and protective variant. Data represent means ± s.d. from three independent experiments. P values shown for one-way ANOVA, and *post hoc* Tukey–Kramer test. * p<0.05, ** p<0.01 and *** p<0.001.

We then examined the effects of the risk variant on expression of a panel of 84 NF-κB target genes. Stimulation resulted in significant upregulation of 23 genes in astrocytes with the protective variant and 28 genes in astrocytes with the risk variant compared to baseline. Transcripts that were differentially expressed by 2-fold or higher (p≤0.05) in astrocytes with the risk compared to the protective variant were IL-15, ICAM1, CXCL10, CCL5 and complement component 3 (C3) (Fig. 2, D, E and S2). Upregulation of protein expression was confirmed with ELISA or Western blot (Fig. 2F and S2). These findings indicate that the rs7665090 risk variant activates a specific set of NF-κB target genes in reactive astrocytes that facilitates lymphocyte recruitment and activation. Moreover, risk variant-induced upregulation of C3 suggests astrocytic polarization towards a recently described toxic phenotype, termed A1 in analogy to M1 macrophages, for which C3 is the main marker^16^.

We subsequently investigated whether the risk variant impacts on homeostatic and metabolic functions of astrocytes, namely glutamate transport and uptake/release of glucose and lactate. Glutamate uptake was robust in unstimulated iPSC-derived astrocytes in both groups and deteriorated with proinflammatory stimulation, as described previously^17^. Removal of glutamate was marginally but significantly more impaired in iPSC-derived astrocytes with the risk compared to astrocytes with the protective variant (Fig. 2G). Glucose uptake did not differ between stimulated and unstimulated astrocytes with either variant, while release of lactate was slightly but significantly diminished in astrocytes with the risk variant after stimulation but not at baseline (Fig. 2, H and I).

### Effect of the rs7665090^G^ variant on hypertrophic astrocytes in white matter MS lesions

Having determined the effect of the rs7665090 risk variant on astrocytes *in vitro*, we asked whether comparable phenotypic changes can be seen in reactive astrocytes in MS lesions. From a total of 14 MS autopsy cases homozygous for either the risk or the protective variant (Fig. S7), we identified 10 cases with chronic active white matter lesions, i.e. lesions with demyelinated lesion centers and lesion edges that contained activated microglia, macrophages and hypertrophic astrocytes (Fig. 3A). For comparison of astrocytic phenotypes associated with the rs7665090 risk and protective variant, we selected lesions with similar CD68^+^ cell densities at the lesion rim (Fig. 3, A and B). Morphological appearance of hypertrophic GFAP^+^ astrocytes at the lesion rim were similar in cases with the risk and protective genotypes. Confocal imaging of immunofluorescent-labeled sections demonstrated universal expression of the NF-κB p50 and p65 subunits in the cytosol and nucleus of hypertrophic astrocytes; however, p50 and p65 was significantly upregulated in astrocytes with the risk variant (Fig. 3, C and D). Presence of NF-κB in the nucleus is an indicator for NF-κB activation, more reliable than p65 phosphorylation, which decays rapidly in *post mortem* tissue^18^. Both p50 and p65 were near absent in non-reactive astrocytes in the lesion vicinity and in normal appearing white matter, indicative of low levels of NF-κB expression in non-activated astrocytes. Enhanced immunoreactivity for NF-κB p50 and p65 in lesions with the risk genotype was not restricted to astrocytes but was also found in GFAP^−^ cells. Moreover, immunofluorescent staining for CXCL10, CCL5, IL-15, ICAM1 and C3d showed substantial upregulation in hypertrophic astrocytes in lesions with the risk variant (Fig. 3, E-G, and S3), as predicted by our results in iPSC-derived astrocytes. As in iPSC-derived astrocytes, a specific set of NF-κB target genes was upregulated, while other markers of astroglial activation (GFAP, iNOS, CCL2 and CXCL1) did not differ between lesions with the risk and protective variant (Fig. 2, D-F and 3G). Moreover, correlation studies revealed strong positive correlations between NF-κB signaling in astrocytes and expression of chemokines, IL15 and C3, while no correlation was found between NF-κB signaling and GFAP/iNOS expression in astrocytes or CD68^+^ cell density at the lesion rim (Fig. S4).

**Figure 3.**
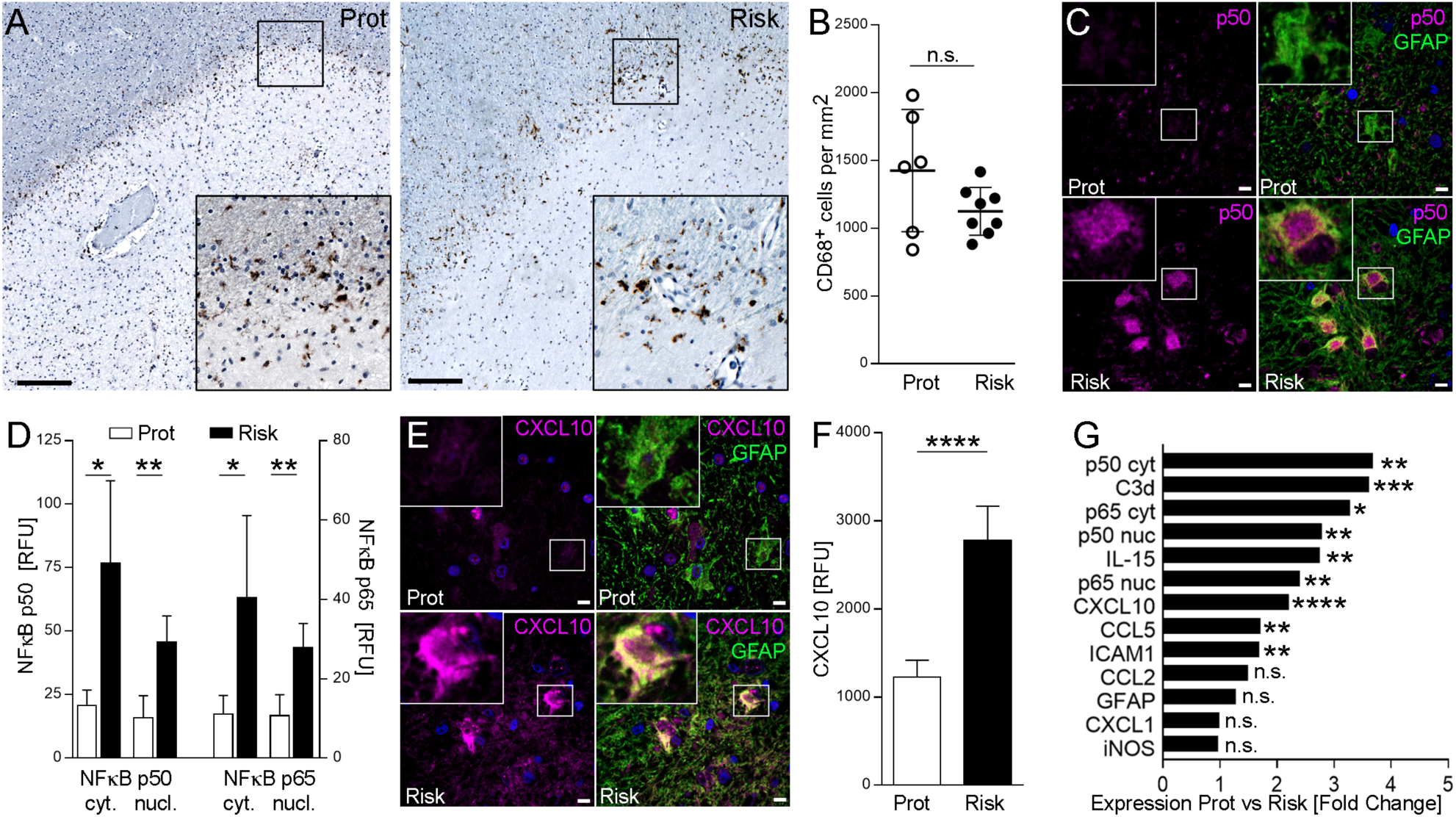
Effect of the rs7665090^G^ variant on hypertrophic astrocytes in white matter MS lesions. (**A**) Chronic active MS lesions from autopsy cases with the rs7665090 risk and protective variant. Staining for CD68 (brown) shows activated microglia at the lesion rim. Counterstain with hematoxylin (blue). Magnifications in inset. (**B**) Quantification of CD68^+^ cell densities in14 lesions from 10 MS cases (five per group). Dots represent average cell densities per lesion counted per group. (**C**) Confocal microscopy images of hypertrophic astrocytes at the lesion edge labeled with fluorescent antibodies against p50 (magenta) and GFAP (green); counterstained with Hoechst 33342. (**D**) Densitometric quantification of NF-κB p50 and p65 in the cytosol/nucleus of hypertrophic lesional astrocytes (5 cases per group). (**E**) Hypertrophic astrocytes stained with fluorescent antibodies against CXCL10 (magenta) and GFAP (green); counterstained with Hoechst 33342. (**F**) Quantification of CXCL10 in hypertrophic GFAP^+^ astrocytes (5 cases per group). (**G**) Differential expression of immune mediators and activation markers in astrocytes from both groups. Data represent means ± s.d. P values shown for one-way ANOVA, and *post hoc* Tukey–Kramer test. * p<0.05, ** p<0.01 and *** p<0.001. Scale bar = 100 µm in A, 15 µm in C, E.

### Effect of the rs7665090^G^ variant on lesion pathology and lesion load in MS patients

Since increased astroglial expression of CXCL10, CCL5 and ICAM1 implies enhanced recruitment of lymphocytes, we quantified number of perivascular lymphocytes within MS lesions as well as lesion size in autopsy cases from both groups. We found that both number of perivascular CD3^+^ T cells, normalized to lesion size (Fig. 4A, B), and lesion sizes (Fig. 4D) were significantly higher in lesions with the risk variant compared to the protective variant, as was the ratio between perivascular CD3^+^ T cells and CD68^+^ microglia at the lesion rim (Fig. 4C). We found strong correlations between CD3^+^ cell infiltration within lesions and astroglial expression of cytosolic p50 (r = 0.80) and CXCL10 (r = 0.67), and between lesion sizes and astroglial expression of C3d (r = 0.81) and IL15 (r = 0.88) (Fig 4E, F; Fig S4).

**Figure 4.**
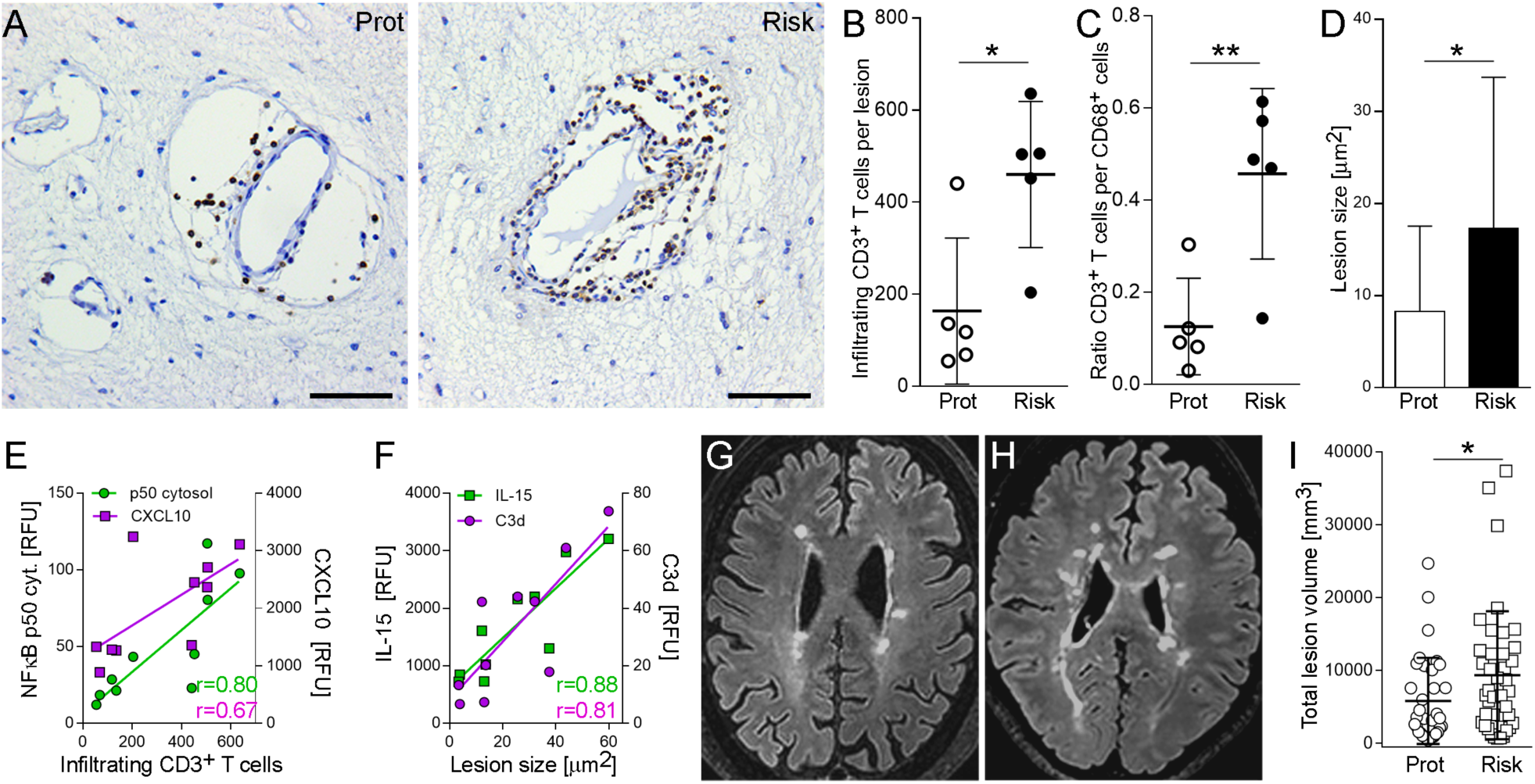
Effect of the rs7665090^G^ variant on lesion pathology and lesion load in MS patients. (**A**) Bright-field images of perivascular infiltrates within chronic active MS lesions labeled with anti-CD3 antibody. (**B**) Quantification of infiltrating, perivascular CD3^+^ T cells in 10 lesions from five MS cases per group. Dots represent average CD3^+^ cells. (**C**) Ratio of CD3^+^ cells per CD68^+^ cells in lesions from both groups. (**D**) Quantification of lesion sizes determined as areas of demyelination in MBP-stained sections (18 lesions from 6 cases (protective) and 21 lesions from 8 cases (risk)). (**E**) Correlation of infiltrating CD3^+^ cells with cytosolic NF-κB p50 (r=0.80, p=0.005) and CXCL10 (r=0.67, p=0.036) expression. (**F**) Correlation of lesion size with IL-15 (r=0.88, p=0.001) and C3d (r=0.81, p= 0.005). **(G, H)** Examples of white matter lesions on FLAIR images from MS patients homozygous for the protective (G) and risk variant (H). **(I)** Total lesion volumes of 35 MS patients (rs7665090^GG^) and 40 patients (rs7665090^AA^) with established disease (≥5 years disease duration). Data represent means ± s.d. p values shown for one-way ANOVA and *post hoc* Tukey–Kramer test (B, C, D) or F-Test (I). * p<0.05 and ** p<0.01. Scale bar=50µm.

Finally, we examined in a cross-sectional study the effect of the rs7665090 risk variant on white matter lesions in MS patients, as measured by MRI. Since individual lesions can become confluent in advanced MS and are thus difficult to delineate, we quantified overall lesion load per patient rather than individual lesion sizes. We recruited a total of 93 patients homozygous for the risk or protective genotype from the MS clinics in Yale, USA and Turku, Finland (50 and 43 patients, respectively). From these, we excluded 18 patients from analysis with < 5 lesions (n=2) and/or < 5 years of disease duration (n=16). In patients that were treated with the highly effective disease modifying therapies that prevent formation of new lesions by >80% (Tysabri, Rituxan, Ocrevus and Lemtrada^19-21^), we defined disease duration from onset of symptoms until the initiation of these treatments, thereby accounting for treatment history. In the remaining 75 patients (patient demographics in table S8), lesion load on fluid-attenuated inversion recovery (FLAIR) images was determined using an automated segmentation algorithm^22^, followed by manual correction by three independent raters. We found that the risk variant was associated with a significant higher lesion load in patients with established disease (Fig. 4G-I). Patient age, gender or disease duration did not significantly affect the white matter lesion load (table S9). Although lesion load is weakly correlated with disease duration^23,24^, this correlation was not seen in our cohorts, presumably because we only included patients with ≥5 years of disease duration.

## Discussion

We examined how a specific risk variant for MS susceptibility, rs7665090^G^, impacts on astrocyte function both *in vitro* and in active MS lesions and on lesion pathology. We demonstrated that the risk variant is associated with enhanced astroglial NF-κB signaling and upregulation of specific NF-κB targets that drive lymphocyte recruitment and neurotoxicity, implied by increased expression of C3d^16^. Homeostatic and metabolic functions of astrocytes were only moderately affected by the risk variant but may contribute to cellular damage in lesions by inducing excitotoxicity and metabolic uncoupling from axons/neurons.

As astrocytes play a major role in controlling leukocyte traffic into the CNS^25^, excessive upregulation of chemokines and adhesion molecules amongst others in astrocytes is likely to increase accessibility for peripheral immune cells to the CNS. This is confirmed in MS lesions where the risk genotype was associated with enhanced lymphocytic infiltration, in addition to larger lesion sizes and higher lesion load on FLAIR images in MS patients. We therefore hypothesize that the risk variant confers MS susceptibility by increasing CNS access to activated lymphocytes, which lowers the threshold for MS lesion formation and ultimately tips the balance towards developing MS. The rs7665090 risk variant is not known to increase MS severity or likelihood of disease progression. Indeed, GWAS of disease severity have to date not yielded candidates at genome-wide significance^26^ and a recent study found that genetic risk variants for MS susceptibility do not influence disease severity^27^, suggesting that rs7665090 risk variant-mediated lymphocyte recruitment does not drive MS severity. Our data show a moderate association between risk variant and increased white matter lesion load in MS patients; however, although counterintuitive, T2 lesion volume does not correlate with disability as measured by EDSS and is not a surrogate marker for disease severity^24,28^.

Moreover, we found that the impact of the rs7665090 risk variant on NF-κB signaling and on NF-κB target gene expression in astrocytes was substantial, consistent with other studies that report large variant effects on individual cellular pathways^5,29,30^. The robust increase in NF-κB activation contrasts with the minor increases in MS susceptibility conferred by the rs7665090 and other risk variants outside of the major histocompatibility complex^6^, indicating that individual pathways, even if strongly dysregulated, play only minor parts for overall MS susceptibility. The effect of the rs7665090 risk variant on lesion load in MS patients was only moderate, presumably because multiple other pathways contribute to lesion formation.

Furthermore, the variant-associated changes in lesion pathology are not likely to be caused by a variant effect exclusive to astrocytes. The rs7665090 risk variant has been shown to enhance NF-κB signaling in lymphocytes^5^ and may modulate NF-κB dependent responses in other cell types relevant to lesion formation, including lymphocytes, macrophages, microglia and endothelial cells. Thus, enhanced lymphocytic infiltration in MS lesions may be caused by enhanced astroglial recruitment but also by increased lymphocyte activation. The inability to discern a variant effect on single cell types in MS lesions is a limitation of our study. However, the rs7665090^G^ variant effect cannot be tested in a mouse model, because the genetic architecture of non-coding regulatory regions of *NFKB1* differs considerably between humans and mice and because the causative variant in the rs7665090-tagged haplotype block that contains over 90 variants is not known, precluding genome editing on a single base level.

Our study suggests that MS risk is driven not only by peripheral immune cells but also by dysfunctional CNS cells. The rs7665090 risk variant may thus contribute to MS risk by promoting exaggerated responses in astrocytes that enhance lymphocyte recruitment and thus direct dysregulated peripheral immune cells to the CNS. In addition to increasing susceptibility for MS, the rs7665090^G^ variant also confers increased risk for primary biliary cholangitis (PBC), an autoimmune disease of the liver associated with lymphocytic infiltration^31^. A potentially important cell type in the pathophysiology of PBC are hepatic stellate cells. These cells exhibit striking morphological and functional similarities to astrocytes, including expression of GFAP, regulation of the blood-tissue barrier and recruitment of phagocytic Kupffer cells in a NF-κB-dependent manner^32,33^. Therefore, the rs7665090 risk variant induced risk for PBC may be driven by excessive NF-κB signaling in stellate cells and stellate cell-mediated recruitment of peripheral immune cells. Conversely, NF-κB related variants located in chromatin regions not accessible in astrocytes (rs1598859 and 3774959) are associated with risk for inflammatory bowel disease and systemic scleroderma but not MS, suggesting that these variants do not impact on lymphocyte recruitment to the CNS. We therefore propose the general concept that autoimmunity is the result of risk variant-driven dysregulation of the peripheral immune system and of cells in the target organ that play a role in locally establishing autoimmune inflammation.

In summary, our study provides evidence for the first time that genetic variants associated with MS risk directly perturb CNS cell functions. It remains to be shown that risk variants impact on other CNS-constituent cells and possible on non-immune immune pathways, that provide plausible mechanisms for increased MS susceptibility. To this end, a systematic correlation of MS risk variants with detailed chromatin landscapes of CNS cells may help delineate the potential relevance of risk variants in different CNS cell types and identify CNS-intrinsic disease-causing pathways. Two recent studies on human microglia found that a majority of candidate genes associated with MS risk alleles were preferentially expressed in microglia^4,34^, further strengthening the idea that CNS-intrinsic mechanisms contribute to MS susceptibility. Presence of genetic variants may have implications for therapeutic approaches in MS patients. A recent study suggested that a specific risk variant, rs1800693, is predictive of adverse effects of TNFα antagonist treatment in MS patients^35^. Similarly, therapeutic targeting of a pathway affected by one or a combination of genetic risk variants may be highly effective in MS patients that carry these variants.

## Materials and Methods

### Study Design

The aim of this study was to establish the effect of the rs7665090 risk variant on (i) astrocyte function *in vitro* and *in situ*, (ii) on MS lesion pathology and (iii) on lesion load in MS patients. IPSC-derived astrocytes, autopsied MS cases and MS patients were all homozygous for either the risk or protective variant. We used astrocyte cultures derived from a total of 18 iPSC line obtained from 6 MS patients. For histological lesion analysis, we examined a total of 39 MS lesions from 14 MS patients. All 39 lesions were used to determine lesion sizes; 10 chronic active lesions were selected for their comparable CD68^+^ cell density and used to determine the degree of CD3 infiltration and astrocyte responses. Finally, we identified 93 MS patients homozygous for the risk or protective variant at the MS Clinics in Yale, USA and Turku, Finland, to determine their lesion load on MRI. From 93 patients we excluded 18 patients from analysis that did not have established disease as defined by either <5 lesions (n=2), <5 years of disease duration or <5 years time to treatment with highly effective disease modifying therapies (Tysabri, Rituxan or Lemtrada; n=16), which prevents formation of new lesions. The primary endpoint and exclusion criteria for MS patients were established prospectively. All outlying data were included in the analysis. Primary data of patients’ characteristics are provided in tables S5, 7 and 8. Sample sizes were dictated by the availability of MS autopsy cases and of genotyped MS patients. In astrocyte culture, we performed at least three independent experimental replicates. From MS autopsy tissue, we selected lesions with comparable CD68^+^ densities at the lesion rim and subsequently determined expression levels from at least 20 reactive astrocytes per lesion. Analysis was performed in all experiments in a blinded fashion.

### Generation of iPSCs from MS patients and differentiation of iPSCs into astrocytes

We obtained skin punch biopsies from six MS patients at the Yale MS Clinic that were homozygous for the rs7665090 risk variant or protective variant (table S5) and generated explant cultures for derivation of primary human fibroblasts as described ^36^. Fibroblasts were cultured by Tempo Bioscience, Inc (San Francisco, CA, USA; http://www.tempobioscience.com/), reprogrammed to three individual iPSC colonies per patient and expanded, using their proprietary reprogramming protocol. Pluripotency of all 18 iPSC lines was confirmed by assessing expression of biomarkers including Oct4, Tra-1-80, Nanog, and SSEA4. IPSC colonies were differentiated into astrocyte progenitors and matured into astrocytes, expressing S100beta and GFAP, using Tempo Bioscience’s serum-, feeder-, integration-and genetic elements-free, non-viral technology. For maturation, astrocytes were plated on poly-L-ornithine/laminin and differentiated with DMEM/F12/neurobasal medium (50%/50%) containing 10ng/ml BMP-4 for 2-4 weeks until >90% of the cells were GFAP^+^ and GLAST^+^ as determined by flow cytometry. FACS-sorted GLAST^+^ astrocytes were used for all experiments.

### Chromatin accessibility profiling in iPSC-derived astrocytes

Human fetal astrocytes (n=2) and iPSC-derived astrocytes (n=4) were profiled for chromatin accessibility using the assay for transposase-accessible chromatin (ATAC-seq) ^12,37^. Aliquots of 5,000 cells were incubated with transposase solution containing 1% digitonin ^37^ at 37°C with agitation at 300 rpm for 30 min. After transposition, DNA was purified (MinElute PCR Purification Kit; QIAGEN) and transposed fragments were minimally PCR amplified ^12^ and purified using the Agencourt AMPure XP system (Beckman Coulter). The average fragment size was estimated by Bioanalyzer (Agilent) and libraries were quantitated with the qPCR-based Library Quantification Kit (KAPA Biosystems). Purified libraries were sequenced on the Illumina HiSeq 2000, generating paired-end 100 bp fragments. Fragments were aligned to hg38 with bowtie2 ^38^ and unique, singly mapping reads were retained for further analysis. Visualization tracks were generated by calculating the number of reads aligning at each genomic position and normalizing for library size.

### Gene expression profiling in iPSC-derived astrocytes

Astrocytes were stimulated with 10 ng/ml IL-1β/100 U/ml IFN-γ for 12 hrs and 50 ng/ml TNF-α for an additional 4 hrs before total RNA extraction (RNeasy Micro Plus Kit; Qiagen), reverse transcription (RT^2^ First Strand Kit, Qiagen) and gene profiling with a Human NF-κB Signaling Targets RT² Profiler PCR Array (Qiagen), as described before ^39^.

### Flow cytometry, Western Blot and ELISA in iPSC-derived astrocytes

To assess NF-κB activation with flow cytometry, astrocytes were stimulated with 50ng/ml TNF-α and 10ng/ml IL-1β for 10 min, dissociated with accutase, stained with fluorescent-labeled primary antibodies (table S6) and analyzed on either a FACSCalibur or a LSRII flow cytometer (BD Biosciences)^39^. Protein expression in unstimulated and stimulated (50ng/ml TNF-α/10ng/ml IL-1β, 48 hrs) cultured astrocytes were determined with Western blot as previously described ^39^ with primary antibodies listed in Fig. S6. Proteins were visualized with an enhanced chemiluminescence using an ImageQuant LAS 4000 camera (GE Healthcare). Densitometry was performed with ImageJ software and values normalized to loading control (β-actin) were used for analyses. Cytokine release was quantified with sandwich ELISAs (DuoSet ELISA for CXCL10, CCL5, IL-6, R&D Systems; complement C3 ELISA; Abcam). Supernatants from unstimulated and stimulated astrocyte cultures were collected, centrifuged and assayed according to the manufacturer’s instructions.

### Metabolic assays in iPSC-derived astrocytes

Glucose uptake was measured in unstimulated and stimulated astrocytes (50ng/ml TNF-α/10 ng/ml IL-1β; 48 hrs) via incorporation of the fluorescent glucose analog 2-NBDG (Thermo Fisher Scientific). 5µM 2-NBDG was added to cells in low glucose medium (1g/L D-Glucose) for 30 min. After washing with HBSS, intracellular fluorescence was determined with an Infinite M1000 fluorescent plate reader (Tecan). Lactate secretion was determined from stimulated and unstimulated cell culture supernatants using an enzymatic assay, which produces a colorimetric (570nm)/fluorometric (λ_ex_=35nm/λ_em_=587nm) product proportional to lactate content. (Sigma Aldrich). To quantify glutamate uptake, we incubated astrocyte cultures with HBSS buffer containing 0.5µM L-glutamate and L-[^3^H] glutamate (1µCi; PerkinElmer) at a 100:1 ratio for 5 min at 37°C. Cells were rapidly moved onto ice, washed twice with ice-cold glutamate-free HBSS buffer and lysed with 0.1N NaOH solution. [^3^H] radioactivity was measured using a scintillation counter and counts were normalized to total protein levels per sample ^40^.

### Genotyping of formalin-fixed autoptic brain tissue

Formalin-fixed CNS tissue of 82 MS patients were obtained from the PI’s CNS bank and the Colorado Brain Bank. Since most cases from the PI’s MS tissue bank came to autopsy in the 1980s/90s, prior to the era of effective MS therapies, the majority of patients did not receive standard disease modifying treatments.

CNS tissue was genotyped by isolating DNA using the DNeasy Blood and Tissue Kit and QIAamp DNA FFPE Tissue Kit (both Qiagen). Pre-amplification of DNA and genotyping for rs7665090 was performed with the Taqman PreAmp Master Mix Kit and a Taqman genotyping assay (Applied Biosystems). Genotyping was carried out in duplicates and repeated at least three times in separately obtained DNA samples. We identified 8 cases with the rs7665090 risk genotype and 6 cases with the protective genotype (details in table S7).

### Bright-field immunohistochemistry in MS lesions and lesion classification

For lesion characterization, tissue blocks containing white matter lesions, were sectioned, quenched with 0.03% hydrogen peroxide, incubated with primary antibodies against MBP (myelin), CD68 (myeloid cells) and CD3 (lymphocytes), processed with the appropriate biotinylated secondary antibody and avidin/biotin staining kit with diaminobenzidine as chromogen (Vector ABC Elite Kit, DAB Kit, Vector Laboratories), and counterstained with hematoxylin^39^. Controls included isotype antibodies for each primary antibody. White matter lesions were categorized as acute, chronic active and chronic silent. Sizes of all white matter lesion (n=39) were quantified from all 14 cases. In chronic active lesions, we quantified the density of CD68^+^ cells at the lesion rim and selected 10 chronic active lesions (five cases per group) with comparable CD68 densities (Fig. 3A). In these lesions, we counted CD3^+^ lymphocytes within perivascular infiltrates throughout the lesion area (Fig. 4A, B) and examined protein expression in astrocytes (Fig. 3C-G).

### Immunofluorescence in MS lesions

To visualize protein expression in reactive astrocytes within lesions, sections were incubated with primary antibodies, listed in table S6, overnight at 4°C, processed with HRP-conjugated secondary antibodies for 2hrs at RT and reacted with Alexa Fluor tyramide (Life Technologies) for 10 min. Subsequently, sections were dyed with 0.7% Sudan Black and CuSO_4_ to quench auto-fluorescence and counterstained with DAPI. Sections were examined and images acquired on an UltraVIEW VoX (Perkin Elmer) spinning disc confocal Nikon Ti-E Eclipse microscope using the Volocity 6.3 software (Improvision). Images were processed with the ImageJ software ^41^. Cytosolic and nuclear expression of NF-κB p50 and p65 and chemokine expression levels of CCL2, CCL5, CXCL1, CXCL10, as well as ICAM1, IL-15, complement factor C3, iNOS and GFAP were quantified by densitometric analysis of fluorescent immunoreactivity of GFAP-positive hypertrophic astrocytes at the lesion rim. Acquisition and analysis were performed in a blinded manner.

### MRI acquisition and T2 FLAIR lesion segmentation

MRI scans were acquired on a 3T Siemens Skyra scanner (Yale) and 3T Philipps Ingenuity TF PET/MR scanner or a Philips Gyroscan Intera 1.5 T Nova Dual scanner (Turku). Automated lesion detection was performed on T2 weighted fluid attenuated inversion recovery (FLAIR) sequences with the LPA algorithm available as part of the lesion segmentation toolbox (LST) (http://www.applied-statistics.de/lst.htm) *(21)*, followed by manual correction using itk-SNAP software version 3.x by three reviewers blinded to the patients’ genotype.

### Statistical analysis

Data represent means ± standard deviation from three independent experiments. Group comparisons of two samples were carried out by *unpaired student’s t-tests*. Comparisons of up to four groups were analyzed by *one-way ANOVA* followed by the Tukey-Kramer multiple comparison *test.* For multiple comparisons, p values were adjusted for False Discovery Rate. For correlation analysis Pearson correlation coefficients were computed. All values passed the D’Agostino-Pearson omnibus normality test for Gaussian distribution. * p<0.05, ** p<0.01, *** p<0.001 and **** p<0.0001. The effect of the risk variant on total lesion load was assessed by applying a logarithmic transformation in order to better approximate a normal distribution of lesion load values ^42^. To account for confounding factors, we fitted a multivariate linear regression model log (lesion load) that included age, gender, disease duration, and genetic variant. The effects of each factor were assessed by F test (ANOVA) (table S9).

## Acknowledgments

D.P. is supported by the National Multiple Sclerosis Society (RG-1610-26049), by the National Institutes of Health Grant R01 NS102267 and by a generous gift from Stanley Trottman. D.A.H. is supported by National Institutes of Health Grants P01 AI045757, U19 AI046130, U19 AI070352, and P01 AI039671, and the Nancy Taylor Foundation for Chronic Diseases. M.L. is supported by an endMS Postdoctoral Fellowship Award from the Multiple Sclerosis Society of Canada. T.S. is supported by the Banyu Fellowship Program and Uehara Research Fellowship Program. L.A. is supported by the Finnish Academy, the Sigrid Juselius Foundation and a Grant for Multiple Sclerosis Innovation by MerckSerono. T.D.N is supported by the National Multiple Sclerosis Society grant RG-1602-07671 and the National Institutes of Health grant R01 NS090464.

Sequencing service was conducted at Yale Stem Cell Center Genomics Core facility which was supported by the Connecticut Regenerative Medicine Research Fund and the Li Ka Shing Foundation. We would like to acknowledge the Rocky Mountain Multiple Sclerosis Center Tissue Bank for contributing post mortem MS brain tissue for this study. This investigation was supported in part by a grant from the National Multiple Sclerosis Society (RG-1610-26049).

